# SOCfinder: a genomic tool for identifying cooperative genes in bacteria

**DOI:** 10.1101/2023.10.16.562460

**Authors:** Laurence J. Belcher, Anna E. Dewar, Chunhui Hao, Zohar Katz, Melanie Ghoul, Stuart A. West

## Abstract

Bacteria cooperate by working collaboratively to defend their colonies, share nutrients, and resist antibiotics. Nevertheless, our understanding of these remarkable behaviours primarily comes from studying a few well-characterized species. Consequently, there is a significant gap in our understanding of microbial cooperation, particularly in natural environments. To address this gap, we can use bioinformatic tools to identify cooperative traits and their underlying genes across diverse species. Existing tools address this challenge through two approaches. One approach is to identify genes that encode extracellular proteins, which can provide benefits to neighbouring cells. An alternative approach is to predict gene function using annotation tools. However, these tools have several limitations. Not all extracellular proteins are cooperative, and not all cooperative behaviours are controlled by extracellular proteins. Furthermore, existing functional annotation methods frequently miss known cooperative genes. Here, we introduce SOCfinder as a new tool to find cooperative genes in bacterial genomes. SOCfinder combines information from several methods, considering if a gene is likely to (1) code for an extracellular protein, (2) have a cooperative functional annotation, or (3) be part of the biosynthesis of a cooperative secondary metabolite. We use data on two extensively-studied species (*P. aeruginosa* & *B. subtilis*) to show that SOCfinder is better at finding known cooperative genes than existing tools. We also use theory from population genetics to identify a signature of kin selection in SOCfinder cooperative genes, which is lacking in genes identified by existing tools. SOCfinder opens up a number of exciting directions for future research, and is available to download from https://github.com/lauriebelch/SOCfinder.

**Data Summary:** All code and associated files are available at https://github.com/lauriebelch/SOCfinder.

**Impact Statement:** Bacteria cooperate by secreting many molecules outside the cell, where they can provide benefits to other cells. While we know much about how bacteria cooperate in the lab, we know much less about bacterial cooperation in nature. Is cooperation equally important in all species? Are all cooperations equally vulnerable to cheating? To answer these questions, we need a way of identifying cooperative genes across a wide range of genomes. Here, we provide such a method – which we name SOCfinder. SOCfinder allows users to find cooperative genes in any bacterial genome. SOCfinder opens up a number of exciting directions for future research. It will allow detailed studies of non-model species, as well as broad comparative studies across species. These studies will allow cooperation in the wild to be studied in new ways.

## Introduction

The last twenty years has seen a revolution in our understanding of microbial sociality. We have moved from thinking that bacteria and other microbes live relatively independent unicellular lives, to discovering that they cooperate and communicate to perform a stunning array of social behaviours (1–6). This revolution has been largely driven by laboratory-based experiments in a small number of model species, especially *Pseudomonas aeruginosa, Escherichia coli,* and *Bacillus subtilis* (7–11) (Supplement S1). In contrast, we know little about social behaviours in natural populations outside of model species, and we don’t know how the importance of cooperation varies across populations and species. For example, we know that division of labour underpins *Bacillus subtilis* cooperation (12, 13), but we don’t know whether this is true in other species. We know that cheating is important in *Pseudomonas aeruginosa* iron-scavenging (6, 14–16), but we don’t know why it doesn’t appear to be important for the same behaviour in *Burkholderia cenocepacia* (17).

Relatively new genomic approaches offer several ways to study cooperative behaviours in natural populations. These genomic approaches rely on methodologies for identifying genes that control cooperative behaviours. One way to identify such ‘cooperative genes’ is to study the behaviour experimentally, and test whether it is cooperative (18, 19). While these experiments are relatively decisive, they are labour intensive and so not feasible for non-model organisms or large scale across species studies. An alternative approach is to use bioinformatic tools to identify genes for cooperative behaviours (28–33). Comparisons can then be made across species in order to examine how the number or proportion of cooperative genes varies, and if this can be explained by evolutionary theory (20–25). For example, do species where interacting individuals are more likely to be clonally related have more cooperative genes (20)? Alternatively, population genetic approaches can be used to test for ‘signatures’ (footprints) of selection for cooperation, to test if putatively cooperative behaviours really are cooperative in natural populations (26, 27). Other possibilities include comparisons between populations, between species with different lifestyles, or between genes that can undergo different rates of horizontal transfer (25).

The most commonly used bioinformatic tool is PSORTb, which can be used to identify genes that code for extracellular proteins (also known as ‘extracellular genes’) (28). These genes are likely to be cooperative because the proteins can diffuse away from the cell. Any effect of the protein, such as breaking down food or neutralising antibiotics, can therefore provide benefits to the whole group of cells (21–25). Another tool is PANNZER, which predicts the function of any gene based on sequence similarity to known proteins (a process known as ‘functional annotation’) (29). Some functions, like ‘extracellular biofilm matrix’ are known to be cooperative (19).

However, there are several problems with these current methods. First, not all extracellular proteins are cooperative, and not all cooperative behaviours are controlled by extracellular proteins. Some important cooperative behaviours like siderophores are produced by many genes (30), none of which encode extracellular proteins. Second, these methods ignore information about a gene’s location in the genome. Many secondary metabolite genes, including those for siderophores, are clustered together in the genome (30). Functional annotation might label the first and third gene in a cluster as cooperative, but miss the middle gene. Third, existing methods don’t use contextual information on the quality and significance of functional annotation. This can make it difficult to compare across species, as there may be variation in the quality of annotations in different taxa. Fourth, existing methods can be slow to implement on bacterial genomes. Fifth, existing methods don’t account for overlap between methods that are being combined, which can lead to mischaracterization or double-counting of genes.

To address these problems, we provide SOCfinder, a bioinformatics tool to find cooperative genes in bacterial genomes (Figure 1). SOCfinder combines information from several methods, considering if a gene is likely to: (1) code for an extracellular protein; (2) have a cooperative functional annotation; or (3) be part of the biosynthesis of a cooperative secondary metabolite. SOCfinder uses information on the quality and significance of database matches and annotations, and takes around 10 minutes to find cooperative genes in an average bacterial genome on a laptop. A separate list of cooperative genes from each tool is provided as an output, along with a total that avoids double-counting genes. SOCfinder version 1.0 is available as an easy-to-use command line tool, with tutorials, R scripts, and python scripts freely available at github.com/lauriebelch/SOCfinder.

**Figure 1:**
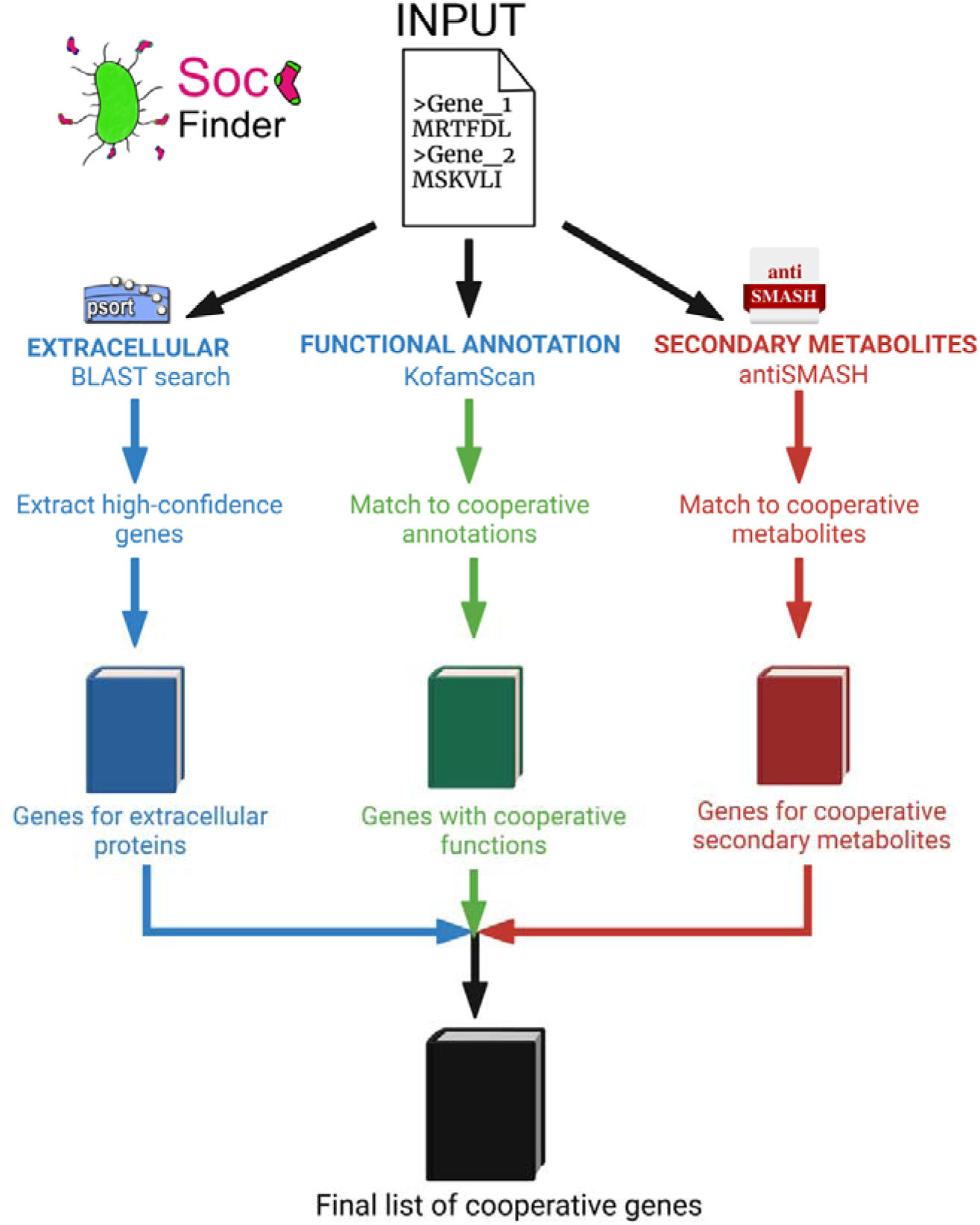
Overview of SOCfinder. We input a genome sequence, and cooperative genes are found based on three modules: (1) Extracellular genes. (2) Genes annotated with functions known to be cooperative, based on sequence similarity. (3) Genes for secondary metabolites that are known to be cooperative. We output a list of cooperative genes for each module, and a final list that combines all three.

We then examine the accuracy of SOCfinder, relative to other bioinformatic tools. We test the ability of different methods to identify genes for cooperation in two species: *Pseudomonas aeruginosa* and *Bacillus subtilis*. We focus on these two species because laboratory experiments have been used to identify a number of cooperative behaviours, including the production of iron scavenging siderophores, quorum sensing and biofilm matrix proteins (7, 31–33). This allows us to test the accuracy and power of the different bioinformatic tools against direct experimental tests. We also test SOCfinder by applying it to >1000 bacterial genomes from 51 species, to see how cooperative gene repertoires vary among and between-species. Finally, we also carry out a population genetic analysis on the genes for cooperation identified by these different tools. This allows us to compare the power provided by the different methods for detecting signatures of selection.

## Methods

### Defining cooperative genes

Before describing our methodology for identifying cooperative genes, we need to define exactly what kind of genes we are looking for. A behaviour is social if it has fitness consequences for both the actor and the recipient (1, 34). Cooperation is a social behaviour where the recipient receives a benefit, and where the behaviour has been selectively favoured at least partially because of that benefit (35). This definition highlights the evolutionary problem of cooperation. Cooperators pay a cost by helping others, so are potentially vulnerable to cheats who benefit from cooperation without paying the cost (36, 37).

In animals, cooperative behaviours tend to be complex traits controlled by many genes, such as worker ants defending the colony (38), vampire bats sharing food (39), or meerkats helping others to rear young (40). As we move from meerkats to microbes the genetics is often simpler, with behaviours involving the production of molecules by one or few known genes. Bacteria produce a range of these molecules that provide benefits to the local group of cells (public goods), including iron scavenging molecules (41), enzymes to digest proteins (42), and toxins to eliminate competitors (43, 44).

We define a cooperative gene in bacteria as a gene which codes for a behaviour that provides a benefit to other cells, and has evolved at least partially because of this benefit. This can be tested for experimentally, by comparing the relative fitness of strains that do and don’t perform a putatively cooperative behaviour both alone and in a mixed culture (1, 7). This contrasts with a ‘private’ gene, which has fitness consequences only for the individual expressing the gene (Figure 2).

**Figure 2:**
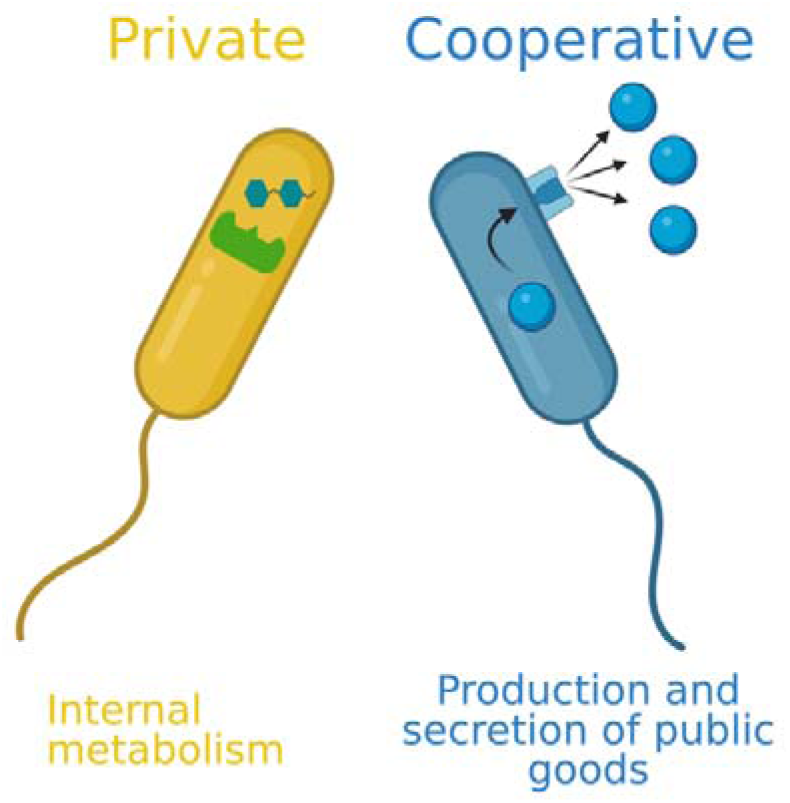
Categorisation of cooperative and private behaviours in bacteria. Cooperative behaviours are involved in the production and secretion of molecules that provide benefits that can be shared with other cells. Private behaviours give fitness benefits only to the individual expressing the gene.

A simple example is *lasB* in the opportunistic pathogen *P. aeruginosa*. This gene codes for the protein elastase, which is secreted outside the cell where it breaks down large structural proteins such as elastin and collagen (45). The digested products can then be taken up by the cell and used for nutrition (46). Lab experiments have compared the growth of the wildtype with a knockout mutant lacking *lasB*. The knockout strain grows slower than the wildtype when grown alone, but outcompetes the wildtype when both are grown together, because it can exploit the elastase produced by the wildtype, while avoiding paying the costs (18, 31, 46, 47). The wildtype is therefore a cooperator and the knockout a ‘cheat’.

### Methods for identifying cooperative genes

In order to assess their validity and usefulness, we examined the methods used by researchers to identify cooperative genes, which vary from simply collating results from experimental work to genome-mining (Figure 3). We examine both the concept behind each method, and the tools used.

**Figure 3:**
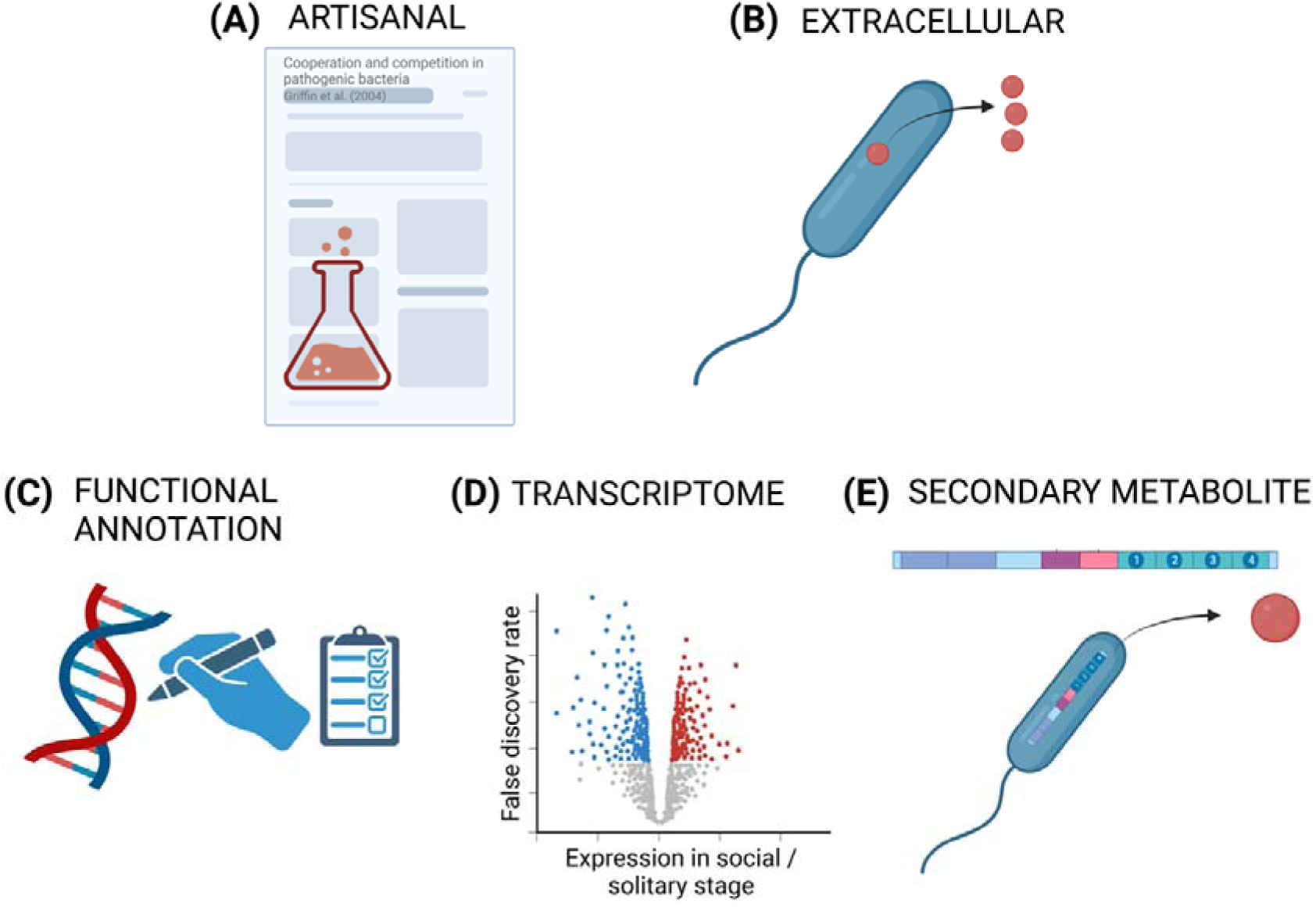
Principles of existing methods to find cooperative genes in genomes. We can look for: (A) Genes that have been shown to be cooperative in lab experiments (**artisan)**. (B) **Extracellular** proteins that are secreted from the cell. (C) Genes that are **annotated** with functions that we know are cooperative, based on sequence similarity to proteins of known function. (D) Genes that are significantly upregulated when individuals are cooperating (**transcriptome**). (E) Genes for the biosynthesis of **secondary metabolites** that are known to be cooperative. A table of specific tools that can be used to find cooperative genes according to these principles is in Supplement S2.

#### Artisanal curation

In some species we can determine the genes for cooperative behaviours, based on upon the results of detailed laboratory experiments. If a species is sufficiently well-studied then we can identify cooperative genes using a literature search for papers conducting these experiments. For example, in *P. aeruginosa*, we could add the gene for elastase *lasB* to our list of cooperative genes based on experimental evidence (18, 31, 46, 47). This method, which we term the ‘Artisanal’ method, has been used in two studies on *P. aeruginosa* (26, 48), and one in *B. subtilis* (27).

#### Extracellular proteins

Many proteins produced by bacteria are extracellular (secreted outside the cell). Genes encoding extracellular proteins are likely to be cooperative because the proteins can diffuse away from the cell and provide a benefit to other cells in the population (22, 25). There are several tools to look for extracellular proteins. For instance, we can use simple BLAST searches to identify extracellular proteins based on similarity to proteins known from lab assays to be secreted, or more sophisticated tools like PSORTb, which also looks at the presence of known sequence motifs (28). This method is the most established for finding cooperative genes, having been used in a number of studies (20–25, 49). One recent study of 51 diverse bacterial species found that on average ∼2% of genes code for extracellular proteins (25).

#### Gene functional annotation

Many gene functions are known to be cooperative, such as the production of extracellular matrix proteins in biofilms. Gene function can be predicted, based on homology and sequence similarity across species for the genes encoding for these behaviours (29, 50, 51). We can use our knowledge of cooperation from model species to make a list of cooperative functional annotation terms, using standardised systems such as gene ontology (GO) or KEGG orthology (KO). For example, Simonet & McNally curated a list of 118 cooperative gene ontology (GO) terms, that can be further split into five categories (secretion systems, siderophores, quorum sensing, biofilm, and antibiotic degradation) (20). They then used PANNZER (29) to predict the function of bacterial genes, which works by looking for homologous sequences which already have GO annotations. Other tools such as KOFAMscan (50) or eggnog-mapper (51) can also be used to predict gene function.

#### PanSort: A combined method

As well as looking at methods in isolation, we can combine the results of multiple methods. This kind of ‘consensus’ method might give better results than any one method in isolation, allowing multiple sources of information to be integrated. This innovative approach was used by Simonet & McNally, who combined a search for extracellular proteins with functional annotation of genes across human microbiome bacteria (20). They used PSORTb to count the number of genes coding for extracellular proteins. They then used PANNZER to annotate gene functions, with the top hit for each gene compared to a curated list of ‘cooperative’ gene ontology (GO) annotation terms. These two totals were then summed to give a total count of the number of cooperative genes in a genome, which could potentially lead to double-counting. We refer to this method, which combined PSORTb and PANNZER, as ‘PanSort’.

#### Transcriptomes

In some microbes there is a distinct social life stage, and we can find the genes controlling this switch in sociality by comparing gene expression between different stages of the life cycle. For example, the bacteria *Myxococcus xanthus* lives in swarms when food is abundant, but upon starvation forms a fruiting body where cells aggregate together. Some cells sacrifice themselves to cooperatively form the stalk that holds up the remaining cells as dispersing spores (52, 53). Similarly, the social amoeba *Dictyostelium discoideum* also has a division between solitary and social life stages (54, 55), with altruistic self-sacrifice in the social stage (56–58). Researchers have used transcriptome data to define cooperative genes as those that are highly expressed in the social stage of the lifecycle, but not in the solitary stage (59).

#### Secondary metabolites

Several known cooperative behaviours in bacteria are not simple extracellular proteins, but are complex molecules developed from several biosynthesis and modification steps. One example is iron-scavenging siderophores such as pyoverdine in *P. aeruginosa* (6, 7, 41). Whilst pyoverdine itself is secreted, none of the proteins controlling its production and export are. Instead, it is a secondary metabolite, defined as a compound that is not required for normal cell growth, but does provide some other benefit (60). We can use bioinformatic tools such as antiSMASH to look for genes that produce secondary metabolites in any genome sequence by looking at sequence similarity and the presence of certain conserved protein domains (61). This tool has been used to help find the cooperative genes that allow *Pseudomonas* and *Paenibacillus* strains to be cooperatively resistant to predation by amoebae when grown together, but susceptible when grown alone (62).

### SOCfinder

Our new method SOCfinder draws on several of these methods. Given an assembled bacterial whole genome, SOCfinder runs three separate modules, and combines the predictions to produce a list of cooperative genes.

#### Module 1: Extracellular proteins

We designed our own method for finding genes that code for extracellular proteins, using the same principles as PSORTb (28). PSORTb gives a prediction of the localization of a protein across the cell, such as the periplasm of cytoplasmic membrane, whereas we only want to know if a protein is secreted or not. We therefore simplified and adapt the BLAST approached used by PSORTb to find gens for extracellular proteins, with some controls to check if a protein matches better to another location. This approach allows SOCfinder to be much quicker than PSORTb.

In our extracellular module, a BLAST search is performed against three out of four custom BLAST databases, based on the subcellular localisation of proteins as determined by PSORTb (Table 1). Depending on whether the species is gram negative or gram positive, either database 1 (gram-positive) or database 2 (gram-negative) is used, whereas databases 3 & 4 are always used (Figure 4).

**Figure 4:**
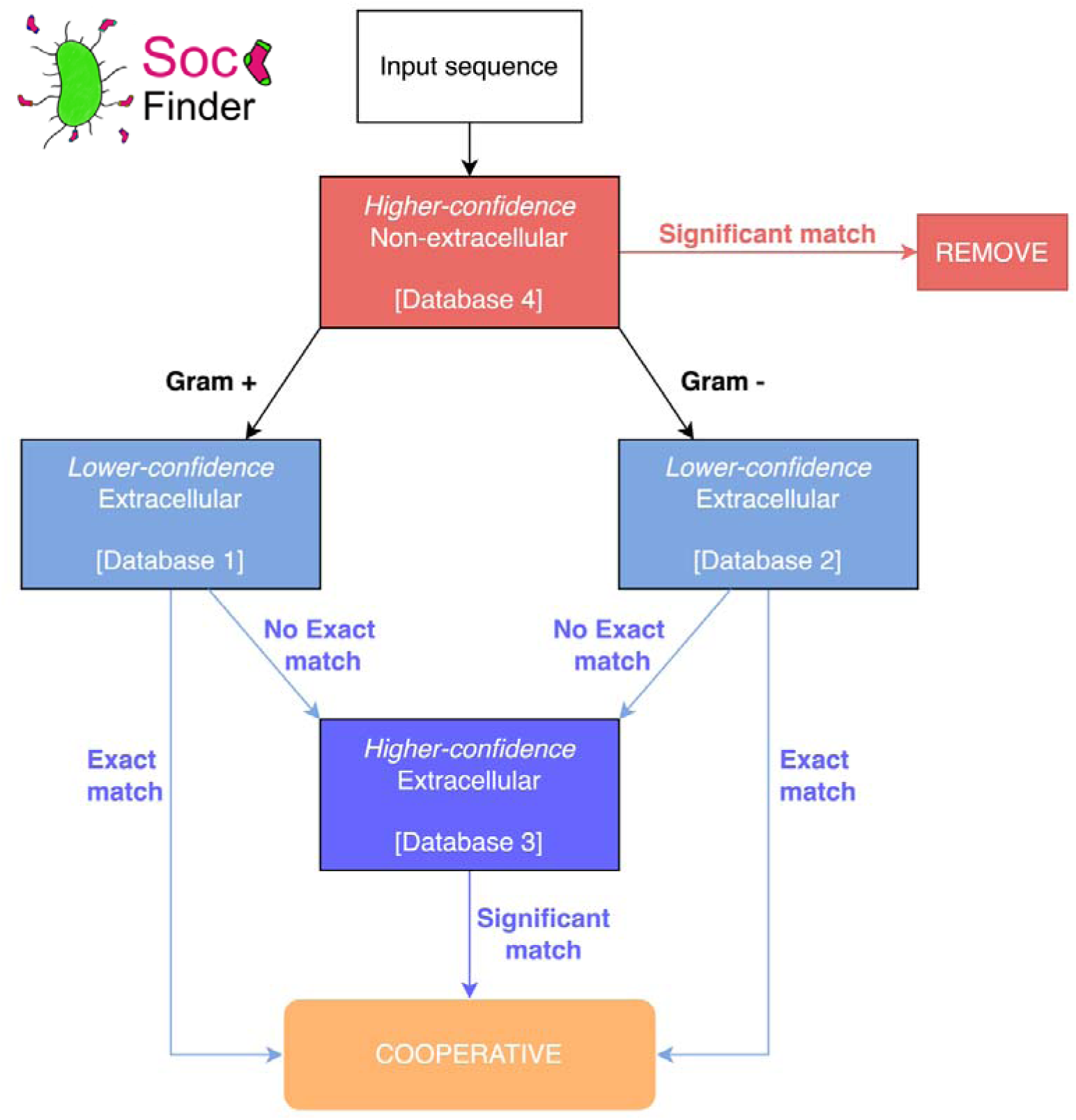
Flow diagram of the BLAST process for finding cooperative genes. Gram-positive and Gram-negative genomes are run against their own databases of high-confidence non-extracellular proteins (database 1 or 2), but both are run against the same databases of higher-and lower-confidence extracellular proteins (databases 3 & 4). Full information of the databases, as well as the definition of a significant match are found in tables 1-3.

**Table 1:** BLAST databases for finding extracellular genes. cPSORT refers to proteins that have been assigned a location based on the PSORTb algorithm. ePSORT refers to proteins with experimental evidence for their localisation.

| Number | Name | Description | Proteins |
| --- | --- | --- | --- |
| 1 | cPSORTdbP<br>extracellular | All the proteins from gram-positive bacteria that are computationally categorised as extracellular by PSORTb3 | 122,392 |
| 2 | cPSORTdbN<br>extracellular | All the proteins from gram-positive bacteria that are computationally categorised as extracellular by PSORTb3 | 156,076 |
| 3 | ePSORTdb<br>extracellular | All the proteins that are categorised as extracellular by the experimentally-derived version of PSORTb4 | 751 |
| 4 | ePSORTdb<br>non-extracellular | All the proteins that are categorised as not extracellular by the experimentally-derived version of PSORTb4 | 9,502 |

**Table 2:** Rules to remove a gene from consideration as cooperative (extracellular). Test Database Action.

| Test | Database | Action |
| --- | --- | --- |
| Query protein has an exact match to a known non-extracellular protein | 4 | Remove from consideration |
| Query protein has a significant* match to a known non-extracellular protein | 4 | Remove from consideration |

\*e-value $<10^{-8}$ , and query and database protein have the same length $\pm 10\%$

**Table 3:**
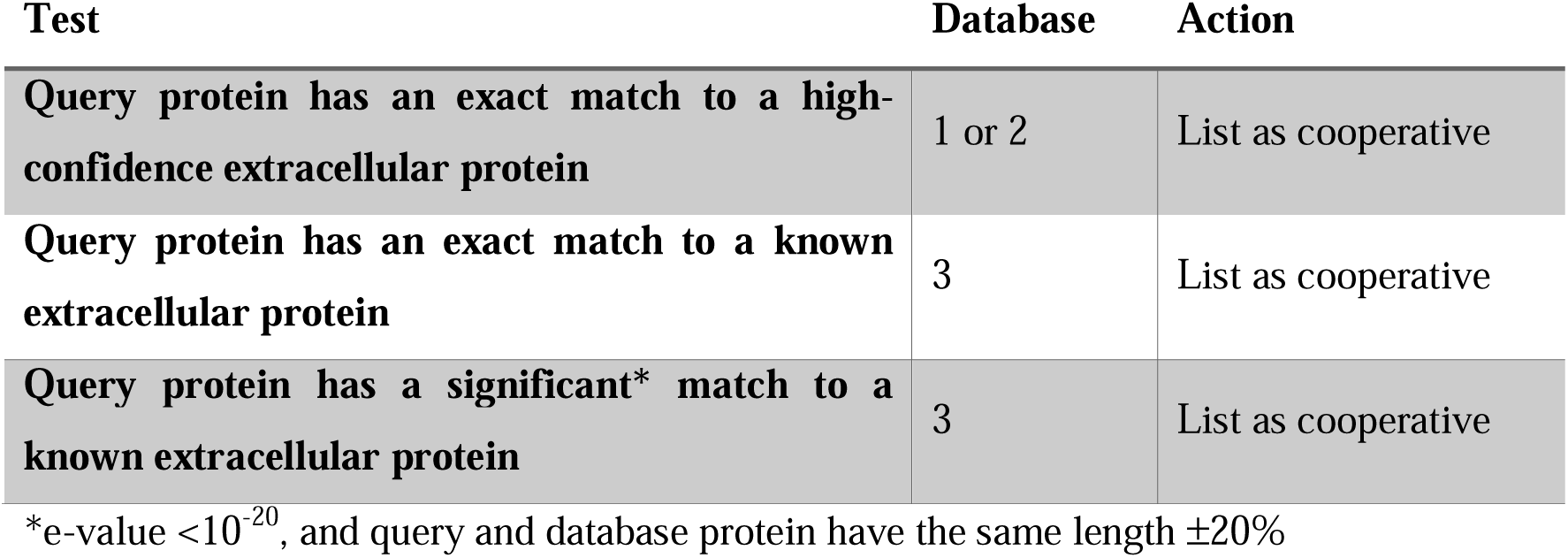
Rules to categorise a gene as cooperative (extracellular).

We first remove some genes from consideration in this module, based on strong evidence that they have a localization that isn’t extracellular (Table 2). This step is important to avoid being too lenient with categorising genes as cooperative. Proteins will often have matches to proteins from multiple localizations, and within a species the same gene can be assigned to different localisations in different strains. We want to have a conservative approach, which is why we apply a stricter significance threshold to include a gene than we do to remove it from consideration, however this can be easily modified by users.

We then test the remaining genes, and categorise genes as cooperative if it meets one or more of the conditions (Table 3). The databases can be found online at https://github.com/lauriebelch/SOCfinder and can be modified by users, and updated as tools such as PSORTb update their own databases to include more genes that have been experimentally or computationally categorised by location.

#### Module 2: Functional annotation

In the functional annotation module, we annotate the genome using KOFAMScan (50). The function of many bacterial genes is known, often because lab experiments have compared the phenotypes of a wildtype and a knock-out mutant that lacks the gene. For any query gene, we can assign it a function based on sequence similarity and machine-learning models that compare our query gene to proteins of known function. The number of matches and the closeness of each match can also be used to assign a score reflecting how confident we are that the query gene really does have that function. The full list of possible functional annotations is held by a database of KEGG orthology (KO) terms, each of which corresponds to a given function (63).

KOFAMScan annotates each protein with any matching KO terms, and each annotation is also given a score as well as an e-value which represents the number of hits it would expect to see by chance for that gene (50). KOFAMScan combines this information to determine whether a given annotation meets its threshold for significance. We can then categorise a gene as cooperative if it has a significant annotation for a KEGG orthology term that is cooperative. To do this, we have created a curated list of cooperative KO terms, generated using a search of all KO terms for keywords corresponding to known cooperative behaviours in bacteria, followed by manual curation to remove KO terms that aren’t likely to be cooperative. The full list of 321 cooperative KO terms is available at https://github.com/lauriebelch/SOCfinder/. Some examples include “exopolysaccharide biosynthesis”, “beta lactamase”, and “pyochelin biosynthesis protein”, and they can be split into nine distinct categories including “siderophore”, “biofilm formation”, and “quorum sensing” (Table 4). For species where we know the full set of genes controlled by quorum sensing, we can use this method to separate cooperative from private quorum sensing genes. Cooperative genes are those highlighted by SOCfinder, and private genes are those not highlighted by SOCfinder. Similar to the extracellular module, we again take a conservative approach. For example, we exclude Type VI secretion systems, which are possibly social (64). However, the user can freely alter this list based on their own criteria.

**Table 4:** Categories of cooperative genes captured by functional annotation.

| Category | Description | Number of KO annotations |
| --- | --- | --- |
| **Beta-lactamase** | Secreted enzymes that provide resistance against beta-lactam antibiotics | 63 |
| **Biofilm formation** | Genes that cause cells to collectively assemble in biofilms | 46 |
| **Exopolysaccharide** | Secreted molecules that form the main part of the biofilm matrix in many species | 56 |
| **Extracellular matrix** | Secreted molecules that form the biofilm matrix | 8 |
| **Quorum sensing** | Genes that regulate or are regulated by quorum sensing, where gene expression changes in response to population density | 88 |
| **Rhamnolipid** | Secreted biosurfactants that allow bacteria to collectively move and disperse over surfaces | 3 |
| **Siderophore** | Secreted molecules that bind to iron, allowing bacteria to scavenge iron from their hosts | 22 |
| **Type II secretion** | Genes secreted by the Type II secretion system used by many gram-negative bacteria to secrete exoproteins into the extracellular environment | 16 |
| **Type IV pili** | Genes for Type IV pili, which are used for collective “twitching motility” | 18 |

#### Module 3: Secondary Metabolites

In the secondary metabolites module, we use antiSMASH (61) to find gene clusters that produce secondary metabolites. The aim here is to ensure that we can capture the entire region for complex social behaviours like iron-scavenging siderophores, where each gene codes for an intracellular protein, but the final product is secreted extracellularly. Functional annotation approaches often capture some, but not all, of these genes. We filter the antiSMASH output to remove all genes which have NA for their ‘type’ (e.g. core biosynthesis, transport, regulation), and then include a gene as cooperative if it matches our custom list of a small number of known social secondary metabolites. Our list includes beta-lactamases and metallophores such as siderophores, which allow bacteria to obtain iron and other metal ions from their hosts (41) (available at https://github.com/lauriebelch/SOCfinder/). Again, this is a conservative approach, but users can easily adjust the list to include other types of secondary metabolite, or as tools such as antiSMASH update their own categorisation.

One of the main strengths of SOCfinder is that it uses three different modules, which tend to capture separate genes. We control for any issues of double-counting by always outputting a final list of cooperative genes that avoids this, whilst still allowing flexibility by outputting separate lists for each module

#### Molecular Population Genetics

We followed the approach used in our previous research of analysing signatures of selection on genes whose expression is controlled by quorum-sensing (26, 27). Population genetic theory predicts that, in non-clonal populations (genetic relatedness *r*<1) that traits favoured by kin selection for cooperation will exhibit increased polymorphism and divergence, relative to traits that provide private benefits (65–69). When comparing putatively cooperative and private traits it is useful to compare traits which are likely to be expressed at similar rates (26, 27). We controlled for expression rates by examining genes controlled by the quorum sensing network. We use published datasets on which genes are controlled by quorum sensing in two species: *P. aeruginosa* and *B. subtilis* (70–73). Within quorum-sensing controlled genes, we assign a gene as ‘cooperative’ if it is found by whichever cooperative method we are testing (SOCfinder, PSORTb, or PanSort). We assign all other quorum-sensing controlled gene as ‘private’.

To analyse a given population genetic measure, we compare three groups of genes: (1) cooperative quorum sensing genes; (2) private quorum sensing genes; and (3) background genes, which are those encoding proteins that localize to the cytoplasm. This set of background genes is least likely to have a cooperative function, and acts as another ’private genes’ comparison.

## Results

### A test of SOCfinder on 51 species

We first tested our method by applying it to 1,301 bacterial genomes from 51 species that were used in a recent study on whether horizontal gene transfer can favour cooperation (25). This allowed us to look at how the number of cooperative genes varies both within-and between species. We found substantial variation across species in the proportion of a genome that is dedicated to cooperative genes, with an average of 2.8% (Figure 5). At one end of the scale, with only 1.2% of its genome dedicated to cooperation is *Buchnera aphidicola,* a symbiont that lives inside aphids (74). At the other end of the scale, with 5.3% its genome dedicated to cooperation is *Chlamydia trachomatis*, an obligate intracellular pathogen (75). Both species have tiny genomes (<1000 proteins), but very different lifestyles. *B. aphidicola* is vertically transmitted and synthesizes amino acids for its host (76). Our estimate here for cooperative genes in *B. aphidicola* is based upon cooperation between bacterial cells, and not cooperative behaviours that it performs to aid its aphid host. However, the search terms in SOCfinder could be expanded to also look at genes for such mutualistic cooperation. *C. trachomatis* has to enter cells, scavenge for nutrients, and fight a hostile immune system – all of which allow lots of opportunity for cooperation (77). Our results also suggest that there can be considerable variation within some species. For example, in *Escherichia coli*, the percentage of cooperation genes varies from 2.3-3.3%, with a median of 2.7%.

**Figure 5:**
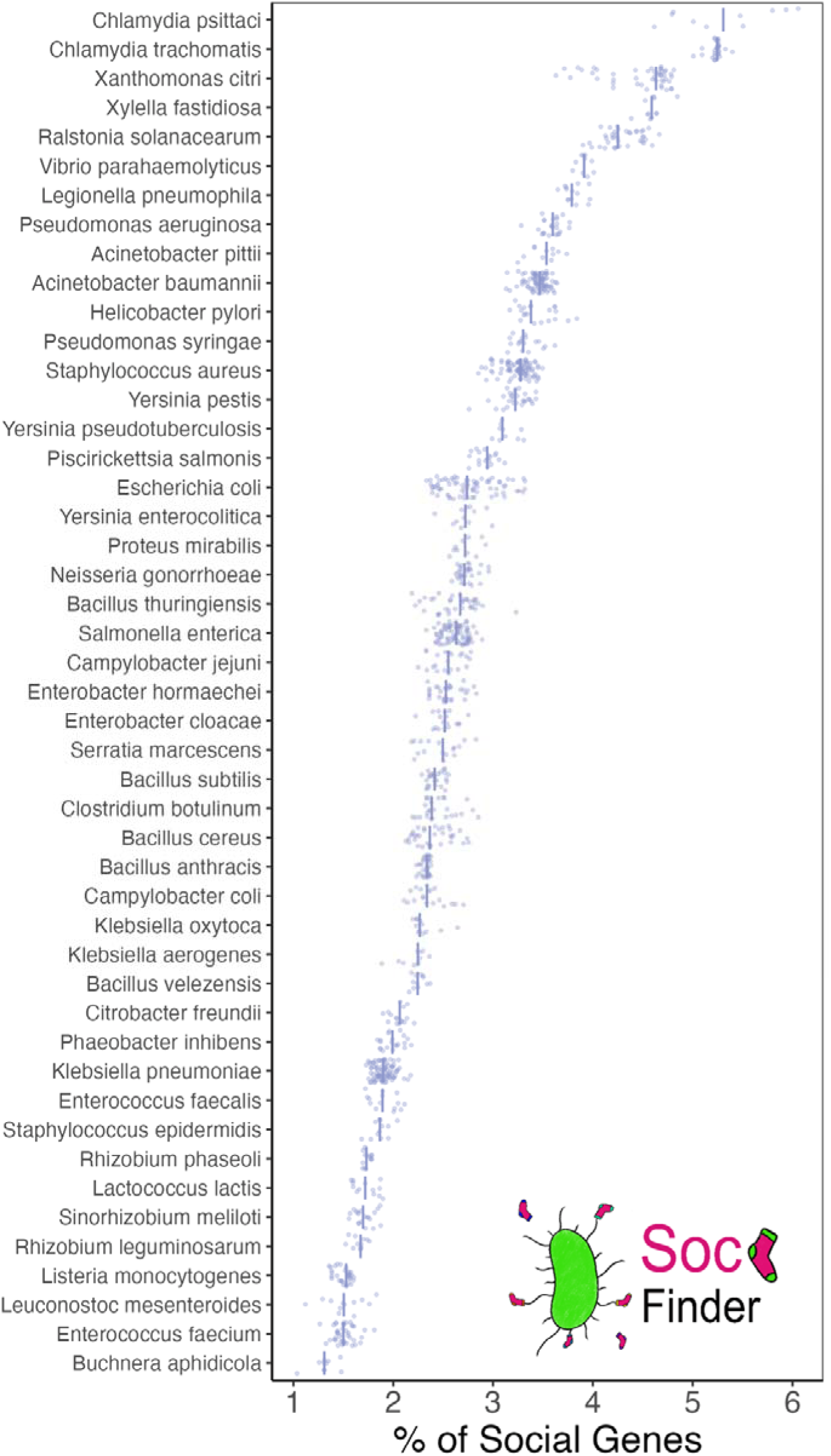
SOCfinder on 1,301 genomes of 51 species. The x-axis shows the proportion of the genes in a genome that are categorised by SOCfinder as cooperative. For each species, a point represents the proportion for one genome, and the bar represents the median proportion.

### Comparison of methods in model species

The artisanal method has been used to identify genes for cooperative behaviours in two well studied species: (1) the gram-negative opportunistic pathogen *Pseudomonas aeruginosa* (26); and (2) the gram-positive soil-dwelling *Bacillus subtilis* (27). In both these species, data from laboratory experiments have identified a number of cooperative behaviours, for which the genes have been determined. We used these artisanal data sets to test the ability and accuracy of other automated methods for identifying genes for cooperative behaviours. We compared three automated methods: (1) the most common previously used method – PSORTb (28); (2) a recent combined method – PanSort (combines PSORTb and PANNZER) (20); and (3) our new method - SOCfinder.

We start by looking at how many genes are captured by each method (Figure 6A&B). SOCfinder captures the most genes. Artisanal captures the fewest genes, because it requires detailed experimental evidence. PanSort and PSORTb are intermediate, with PanSort capturing almost as many genes as SOCfinder, while PSORTb captured many less.

**Figure 6:**
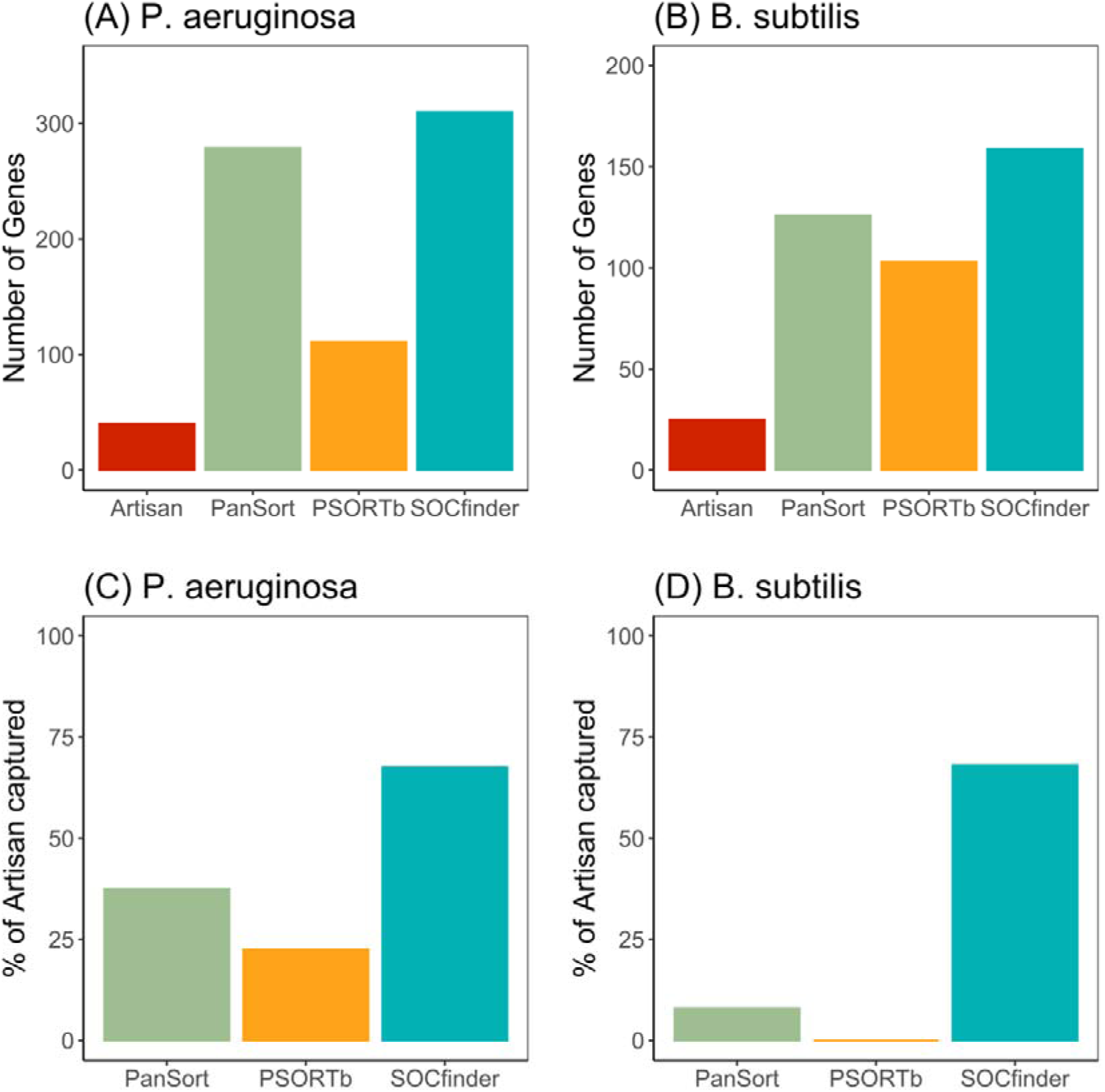
(A&B) Number of genes captured by each method. (C&D) Percentage of artisanal cooperative genes captured by each method. The left panels (A&C) are for *P. aeruginosa*, and the right panels (B&D) are for *B. subtilis*. One key cooperative trait in *P. aeruginosa* is the production of iron scavenging pyoverdine molecules (6, 7, 41). SOCfinder is more than three times better than PanSort at capturing pyoverdine genes, (24/34 = 71%, compared to 7/34 = 21%, binomial test p<10^-9^) (Supplementary Figure 3). PSORTb does not capture any of the pyoverdine genes (Supplementary Figure 4).

We next look at how many of the Artisanal genes are captured by each method (Figure 6C&D). SOCfinder does much better than the other method in both species. In *P. aeruginosa*, SOCfinder captures 68% of the 40 Artisanal genes, which is significantly more than the next best method (38% by PanSort and only 23% by PSORTb, binomial test p<0.001). In *B. subtilis*, SOCfinder captures 68% of the 25 Artisanal genes, which is also significantly more than the next best method (PanSort 8%, PSORTb 0%, binomial test p<10^-12^).

### Can we explain why different methods give different results?

There are a number of possible explanations for the lack of overlap, in terms of genes identified, between the different methods (Figure 7). We now examine the explanatory power of these different explanations, to both test the usefulness of different methods, and guide possible future adjustments to SOCfinder.

**Figure 7:**
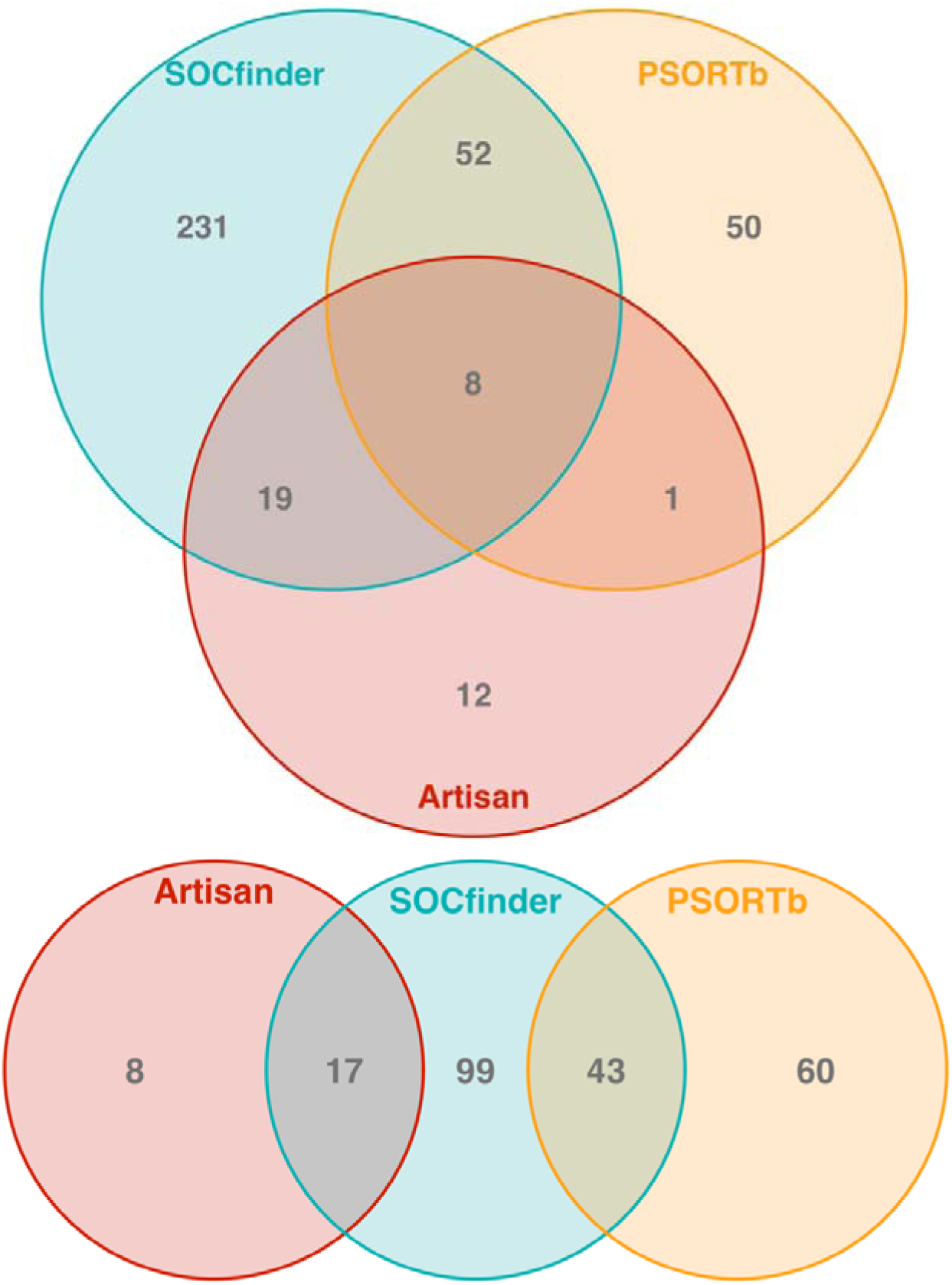
Overlap between methods to find cooperative genes. The top Venn diagram is for *P. aeruginosa*, and the bottom Venn diagram is for *B. subtilis*. The red circle is genes categorised as cooperative by the Artisanal approach. The blue circle is genes categorised as cooperative by SOCfinder. The yellow circle is genes categorised as cooperative (extracellular) by PSORTb.

#### Which known cooperative genes are not found by PSORTb?

There are many known cooperative genes are not extracellular based on PSORTb (31 genes in *P. aeruginosa,* and 25 in *B. subtilis*). Many of these will be intracellular (such as pyoverdine biosynthesis genes), however it is also possible that PSORTb is too conservative in deciding if a gene is extracellular. If this is true, then PSORTB will list the genes as ‘Unknown’ localization (21% of all genes in *P. aeruginosa*, 19% in *B. subtilis*). We tested if the missed cooperative genes are more likely to be listed as ‘unknown’ than the average across the genome. In *P. aeruginosa,* missed cooperative genes aren’t overrepresented for unknown genes (16% of missed genes are unknown, binomial test p=0.66), but in *B. subtilis* they are (32% of missed genes are unknown, binomial test p<0.01).

In gram-negative bacteria which have an outer membrane, another possibility is that PSORTb mistakenly categorises some artisanal cooperative genes as ‘outer membrane’. We tested this in *P. aeruginosa*, and found that cooperative genes missed by PSORTb aren’t overrepresented for ‘outer membrane’ genes (2/31 = 6.5% of missing cooperative genes are outer membrane, compared to 3.1% of all genes: binomial test p=0.244). This isn’t surprising, as we know that many intracellular genes are involved in producing extracellular traits, such as pyoverdine (Supplementary Figures 3&4).

#### Which extracellular genes are missed by SOCfinder?

SOCfinder doesn’t include many genes that are identified by PSORTb as extracellular (51 in *P. aeruginosa*, 55 in *B. subtilis*). This may be because these ‘extracellular but non-cooperative’ genes have no known function, and so wouldn’t have been caught by the functional annotation module of SOCfinder. We found some support for this hypothesis. In both *P. aeruginosa* and *B. subtilis* extracellular genes missed by SOCfinder are significantly more likely to produce a “hypothetical protein” than extracellular genes which are included by SOCfinder (*P. aeruginosa* 21/51 = 41.2% compared to 14/60 = 23.3%, binomial test p<0.01; *B. subtilis* 18/55 = 32.7% compared to 0/48 = 0%, binomial test p<10^-15^).

#### Why are some Artisanal cooperative genes missed by both PanSort and SOCfinder?

There are several known cooperative genes which are missed by both PanSort and SOCfinder (11 genes in *P. aeruginosa,* and 7 in *B. subtilis*). These genes are missed because the annotations they are given don’t match a known cooperative function, although most have a significant annotation (10/12 = 83.3% in *P. aeruginosa;* 3/7 = 42.9% in *B. subtilis*). Often these annotations are too broad to be useful for our purposes, such as “protease I”. Future work is likely to improve functional annotation pipelines, which may allow these missing genes to be eventually captured.

### Can we detect kin selection for cooperation in genes for cooperative behaviours?

Another way to test the usefulness of the different approaches for identifying genes for cooperation is with population genetics. Population genetic theory suggests that selection is relaxed on cooperative genes relative to private genes, making deleterious mutations more likely to fix, and beneficial mutations less likely to fix (65–69). This is because cooperative genes only provide a benefit to carriers of the gene a certain proportion of the time, based on the likelihood that the recipient shares the cooperative gene (genetic relatedness, *r*). Consequently, genes for cooperative behaviours favoured by kin selection, in non-clonal populations (*r*<1) should show increased polymorphism and divergence relative to genes for private behaviours.

Studies on both *P. aeruginosa* and *B. subtilis* have supported this prediction (26, 27). However, these studies used the artisanal approach to identify cooperative and private genes. The artisanal approach was used in these studies because accuracy of identification of cooperative genes is required to be able to pick up possibly subtle population genetic patterns, that could be missed by larger but potentially more messy data sets, compiled with other approaches. In this section, we ask whether other methods to identify cooperative genes give similar results. If the results of an approach do not agree with an analysis on artisanal selected genes, then it could suggest a possible problem with that alternative approach. We examined patterns of polymorphism and divergence for cooperative and private genes identified with three methods: (1) PSORTb; (2) PanSort; and (3) SOCfinder.

When examining genes identified by PSORTb we did not find the expected pattern of increased polymorphism (Figure 8 C&F) and divergence (Supplementary Figures 1&2). There was no significant difference in polymorphism between cooperative and private genes in *P. aeruginosa* (Kruskal–Wallis X2=0.45, p=0.80) or in *B. subtilis* (Kruskal–Wallis X2=2.37, p=0.31). Non-synonymous divergence was significantly higher in cooperative genes compared to private genes in *P. aeruginosa* (Kruskal–Wallis X2=13.2, p<0.01, Dunn Test p=0.03), but not in *B subtilis* (Kruskal–Wallis X2=0.51, p=0.77). Synonymous divergence was not significantly different in cooperative genes compared to private genes in *P. aeruginosa* (Kruskal–Wallis X2=2.86, p=0.24), or in *B. subtilis* (Kruskal–Wallis X2=5.74, p=0.06).

**Figure 8:**
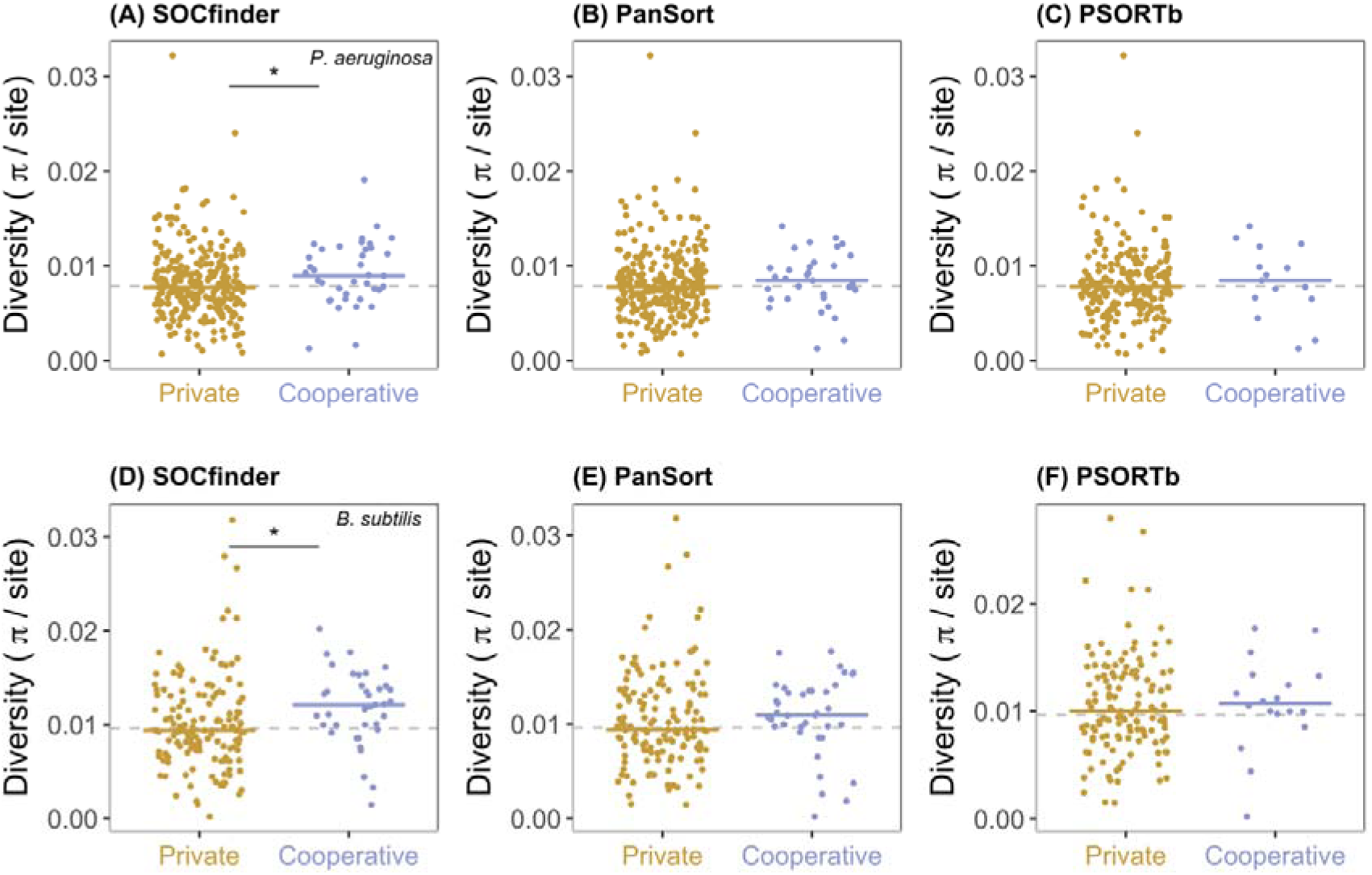
Nucleotide polymorphism for private (gold) and cooperative (blue) quorum-sensing controlled genes. The top three graphs (A-C) show *P. aeruginosa*, and the bottom three graphs (D-F) show *B. subtilis*. The left graphs (A&D) show cooperative genes identified by SOCfinder. The middle graphs (B&E) show cooperative genes identified by PanSort. The right graphs (C&F) show cooperative genes identified by PSORTb. For each graph, the dotted line shows the background level of nucleotide polymorphism for a set of private genes. The black line and * shows a significant difference between cooperative and private genes.

When examining genes identified by PanSort we also did not find the expected pattern of increased polymorphism (Figure 8 B&E) and divergence (Supplementary Figures 1&2). There was no significant difference in polymorphism between cooperative and private genes in *P. aeruginosa* (Kruskal–Wallis X2=1.35, p=0.51) or in *B. subtilis* (Kruskal–Wallis X2=3.81, p=0.15). Non-synonymous divergence was significantly higher in cooperative genes compared to private genes in *P. aeruginosa* (Kruskal–Wallis X2=24.3, p<0.0001, Dunn Test p=0.03), but not in *B subtilis* (Kruskal–Wallis X2=2.28, p=0.32). Synonymous divergence was significantly higher in cooperative genes compared to private genes in *P. aeruginosa* (Kruskal–Wallis X2=9.46, p<0.01, Dunn Test p<0.01), but not in *B subtilis* (Kruskal–Wallis X2=14.73, p<0.001, Dunn Test p=0.26). This indicates that PanSort may be performing better in *P. aeruginosa* than it does in *B. subtilis*.

In contrast, when we identified cooperative and private genes with SOCfinder, we did find that cooperative genes had the signature of kin selection for cooperation, with elevated polymorphism (Figure 8 A&D) and divergence (Supplementary Figures 1&2) compared to private genes. Polymorphism was significantly higher in cooperative genes compared to private genes in both species (*P. aeruginosa*: Kruskal–Wallis X2=6.12, p<0.05, Dunn Test p=0.04. *B. subtilis* Kruskal–Wallis X2=8.48, p<0.02, Dunn Test p=0.01). Non-synonymous divergence was significantly higher in cooperative genes compared to private genes in both species (*P. aeruginosa*: Kruskal–Wallis X2=21.1, p<0.0001, Dunn Test p=0.006. *B. subtilis* Kruskal–Wallis X2=8.26, p<0.02, Dunn Test p=0.02). Synonymous divergence was significantly higher in cooperative genes compared to private genes in *P. aeruginosa* (Kruskal–Wallis X2=9.60, p<0.01, Dunn Test p<0.01), but the trend was significant in *B. subtilis* (Kruskal–Wallis X2=16.70, p<0.001, Dunn Test p=0.08).

## Discussion

We have developed a bioinformatic tool for identifying genes for cooperative behaviours in bacteria. SOCfinder combines information from several methods, and still only takes less than 10 minutes to identify cooperative genes in an average bacterial genome (Supplement S3). Our analyses suggest that SOCfinder both identifies cooperative genes more accurately, and finds more cooperative genes, compared with previous methods such as PSORTb or a combination of PSORTb with functional annotation (PanSort). In addition, these other methods appear to mis-assign genes, to the extent that they are unable to capture the underlying population genetic processes.

The different methods for identifying cooperative genes each have different pros and cons (Table 5). The artisanal method, based on the results of examining behaviours with laboratory experiments represents the relative gold standard in terms of accuracy. It is for this reason that we used it previously when carrying out population genetic analyses, where any incorrect assignments would have introduced noise that could have concealed underlying patterns (26, 27). However, this approach is labour intensive, produces a limited number of genes, and is restricted to species where there has been considerable experimental work, such as *P. aeruginosa* and *B. subtilis*. For example, it identified 40 genes for cooperation in *P. aeruginosa* and 25 genes in *B. subtilis.* Consequently, this approach cannot be applied across the whole genome, to a wide range of species, or to facilitate broad comparative studies.

Methods such as PSORTb are potentially less accurate, but can be automated, and applied across the whole genome of a wide range of species. PSORTb has been used to identify genes for cooperation in a number of studies, for both studies of single species, and broad across species studies (21–23, 25). This has allowed many more genes and many more species to be analysed in a single study. However, PSORTb introduces some inaccuracies with how it identifies cooperative genes, capturing none of the artisanal identified cooperative genes in *B. subtilis*, and only 23% in *P. aeruginosa.* In addition, our population genetic analyses that the level of inaccuracy is sufficient that the noise introduced prevents us from observing the signature (footprint) of kin selection for cooperation at the genomic level.

The importance of the potential problems with using PSORTb can depend upon the kind of question being asked. For example, if you want to know if cooperative genes evolve fast in symbionts, then you need to categorise (‘bin’) genes as either cooperative or private. You don’t want to miss many cooperative genes, because they would then be categorised as private and introduce noise to any comparison. PSORTb could be a problematic approach for such questions. In contrast, if you just wanted to know which intracellular pathogens have the most cooperative genes (‘counting’), then it is less important if you miss some cooperative behaviours. Extracellular genes are likely to be a good proxy for this, and so using PSORTb could be less problematic. The PanSort method developed by Simonet & McNally fixes some of the problems of PSORTb by including some functional annotation (20). However, we show that PanSort doesn’t make full use of power of functional annotation, and still performs badly on the best studied cooperative traits like pyoverdine (Supplementary Figures 3&4), and when comparing to the gold-standard artisanal method.

SOCfinder allows large scale analyses, across whole genomes, and across a broad range of species, but without the same level of problems introduced by PSORTb. SOCfinder is more accurate in identifying cooperative genes because it uses contextual information on the quality of functional annotations, and includes antiSMASH to capture full clusters of biosynthetic genes for key cooperative traits like pyoverdine (Supplementary Figures 3&4). SOCfinder captures variation in the cooperative gene repertoire of bacteria. SOCfinder performs better than other methods in replicating the signature of kin selection that we know exists from studies that have used the gold-standard artisanal approach. To a large extent therefore, SOCfinder has the advantages of methods such as PSORTb, while significantly reducing the disadvantages.

**Table 5:**
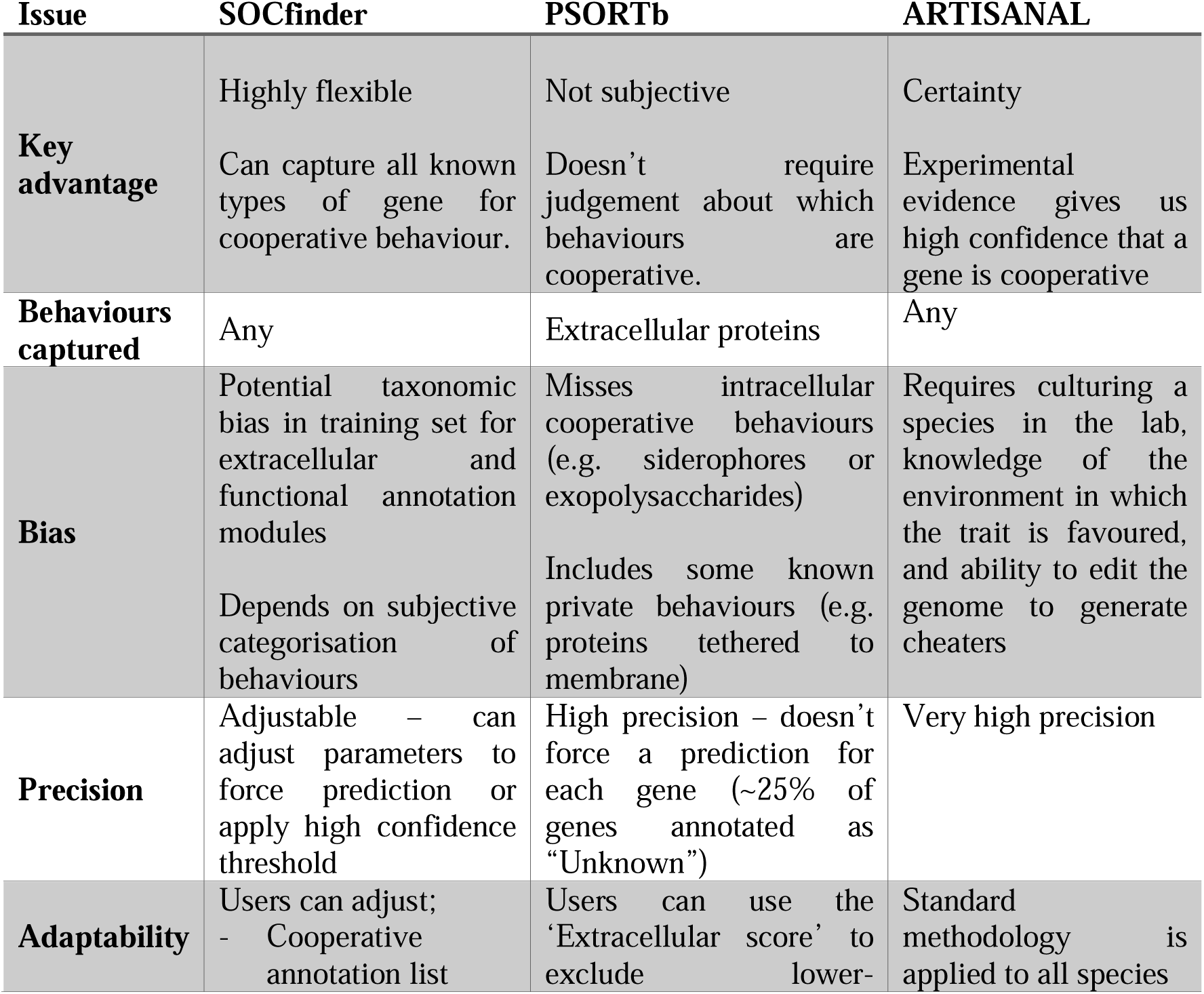
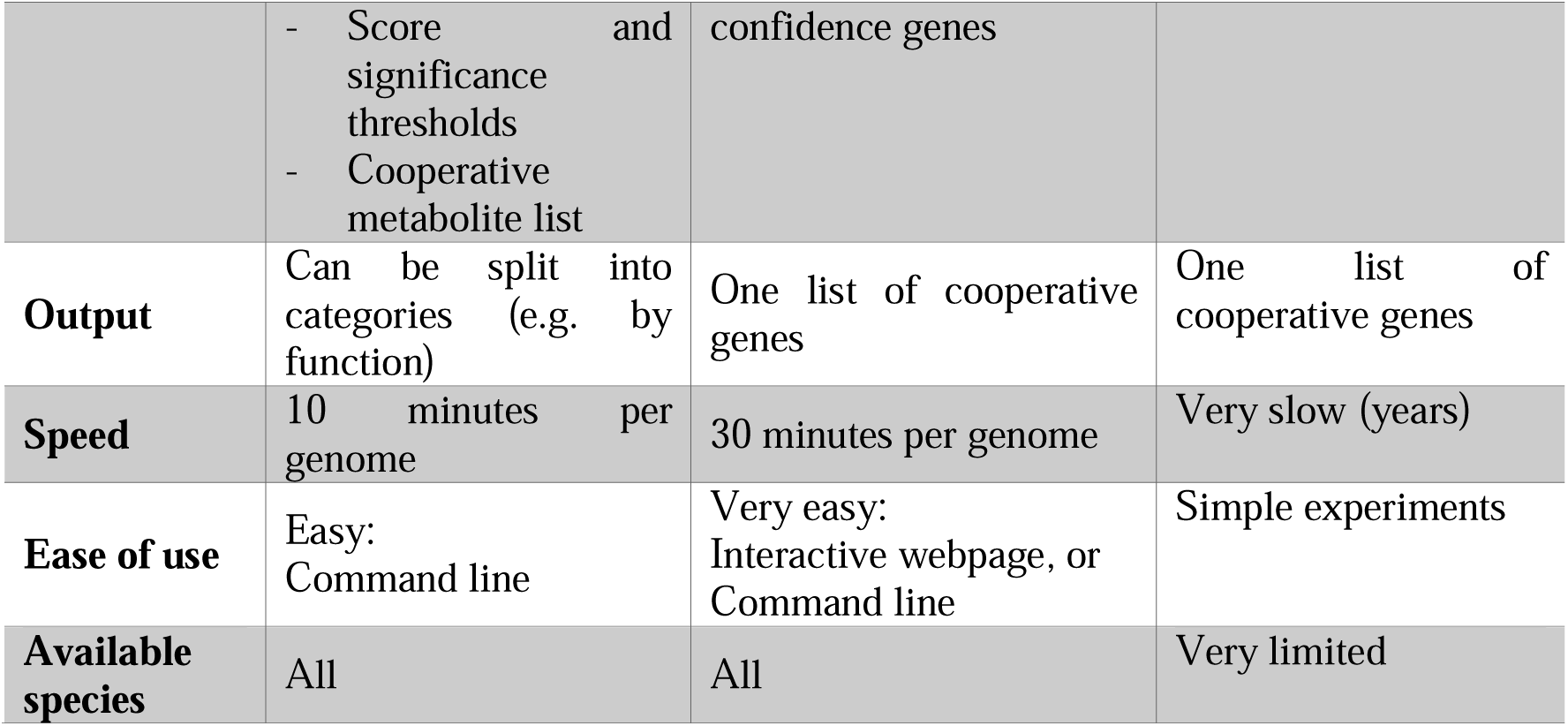
Advantages and disadvantages of methods.

To conclude, SOCfinder opens up a number of exciting directions for future research. It will allow both detailed studies of non-model species, and broad across species studies. These studies will allow cooperation, and how cooperation shapes the genome, to be studied in new ways, such as in natural populations of bacteria. As one example, we could investigate if species that use greenbeards (37, 78, 79) or genetic kin recognition mechanisms (2, 80, 81) have more cooperative genes than those that use environmental kin recognition. In addition, SOCfinder could be used to reassess the results of previous studies which used methods such as PSORTb. We have shown how such methods could lead to limited or inaccurate identification of gene function, and that this could be particularly important if ‘binning’ approaches were used to compare ‘cooperative’ to ‘non-cooperative’ genes. It is still unknown whether the unavoidable inaccuracies imposed by methodologies such as PSORTb have led to biased conclusions.

## Acknowledgements

We thank Carolin Kobras & Ming Liu for useful discussion and comments on the manuscript, and Sarah Flint and Harvey Jeffrey for testing the tool. This work was supported by the European Research Council (834164: LJB, AED & SAW; SESE: MG), and the Biotechnology and Biological Sciences Research Council (BBSRC Oxford Interdisciplinary Bioscience Doctoral Training Partnership ZK). The authors would like to acknowledge the use of the University of Oxford Advanced Research Computing (ARC) facility in carrying out this work.

## Supplement S1: Taxonomic bias

We conducted a Web of Science search for papers on cooperation in bacteria. Specifically, we searched for [Topic = “cooperation” OR “public good” AND “bacteria”] AND [Year = since 2000] AND [Type = “article”] AND [Category = “microbiology” or “evolutionary biology”]. This gave n=464 papers.

We took the list of bacteria genera from the Approved List of Bacterial Names (https://lpsn.dsmz.de/). We included only those genera which are culturable, validly published, and have a correct name (*n*=4180 genera).

We then looked for genus names in the title, keywords, and abstract of the papers. 308 of the papers have a genus name mentioned in one of those three places. More than 28% of these papers mention *Pseudomonas*, with even *Escherichia* and *Bacillus* lagging behind (Figure S1.1)

**Figure S1.1.**
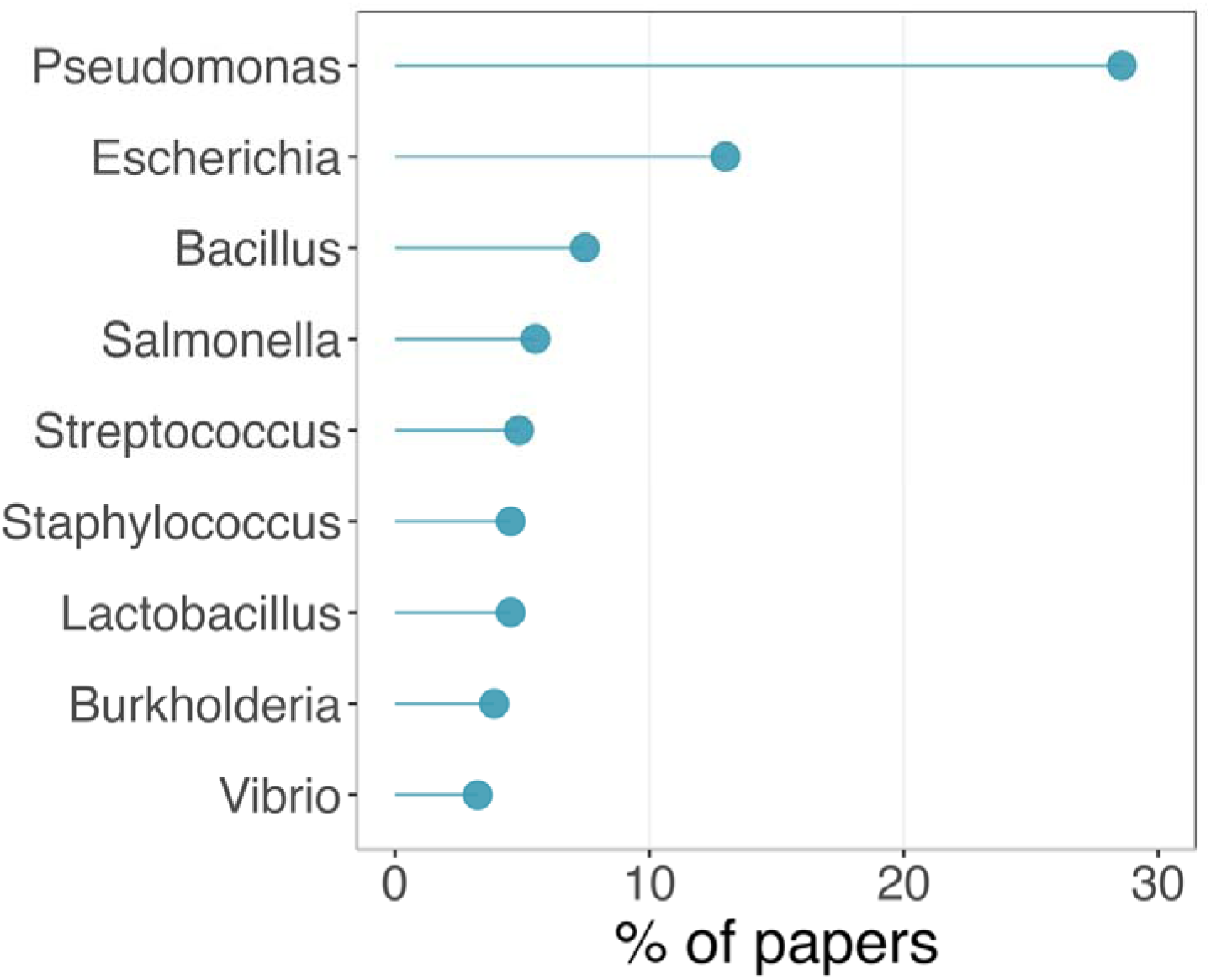
The percentage of papers about microbial cooperation that mention each genus in the title, abstract, or keywords.

## Supplement S2: Other tools

**Table S2:**
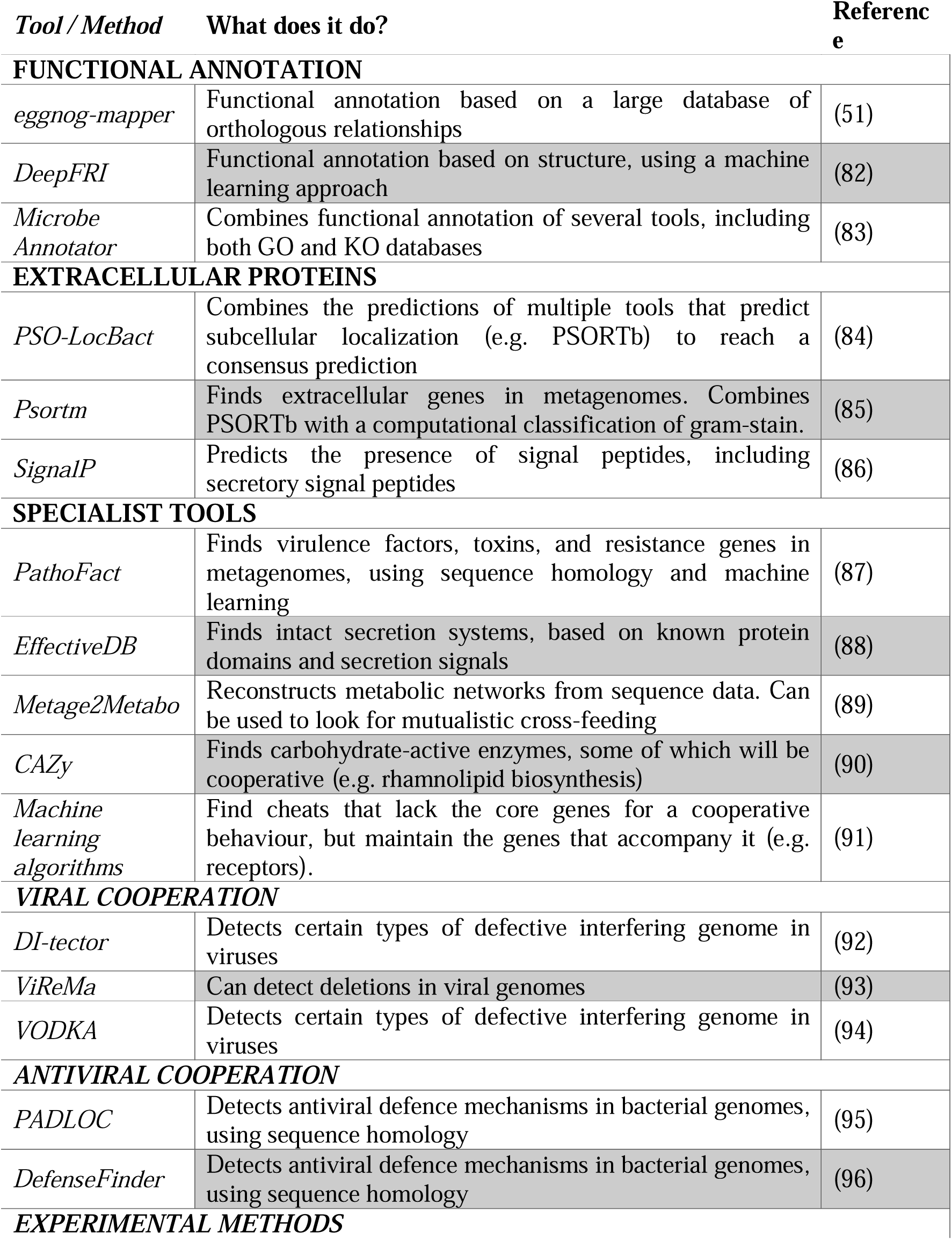
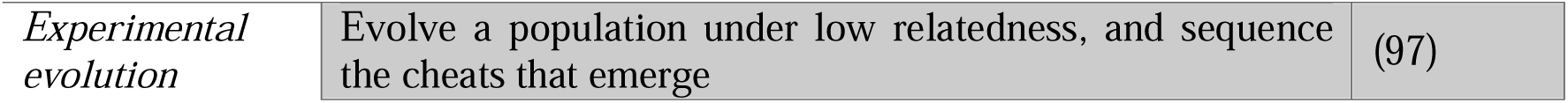
List of alternative tools and methods to find cooperative genes. Tools are separated based on what they find, how they find it, and what species they work on.

### Supplement S3: Speed test

We ran a small test to compare the speed of SOCfinder with the speed of PanSort. PSORTb is the slowest part of PanSort, so the time for PanSort is equivalent to the time for PSORTb.

We chose ten *Escherichia coli* genomes for our speed test (accession numbers GCA_013357365.1, GCA_005221885.1, GCA_004358365.1, GCA_030013595.1, GCA_008931135.1, GCA_009650035.1, GCA_006874785.1, GCA_001612475.1, GCA_024223415.1, GCA_024223415.1). The test was run on a 2020 iMac with a 3.8GHz 8-core intel i7 processor and 32GB RAM.

SOCfinder is significantly quicker than PanSort (t-test, t=109.95, df=17, p<10^-15^), taking 8 minutes on average per genome, compared to 32 minutes for PanSort (Figure S3.1).

**Figure S3.1.**
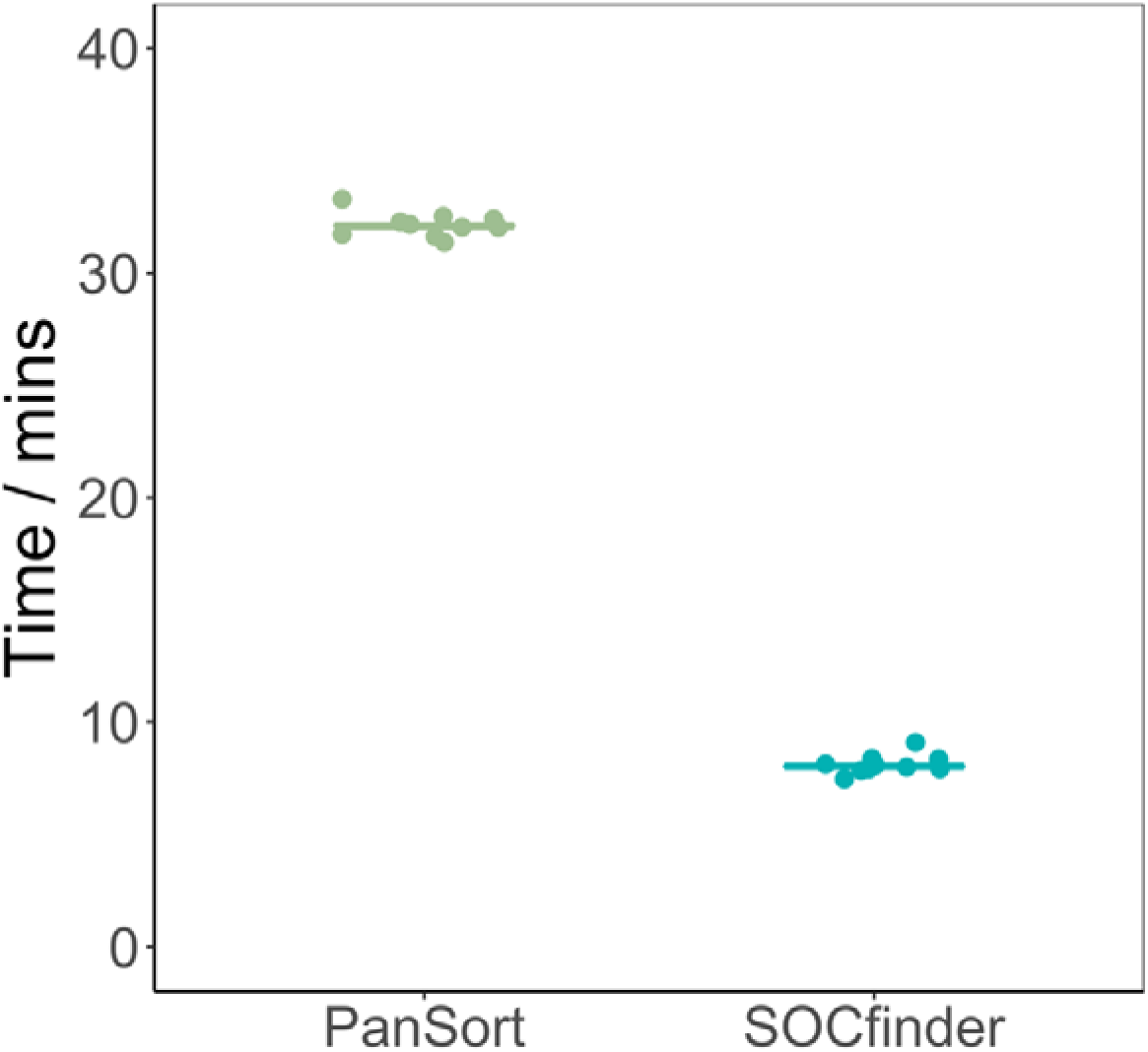
Time in minutes for PanSort (green) and SOCfinder (blue) to run on ten *E. coli* genomes.

### Supplementary Figures

**Supplementary Figure 1:**
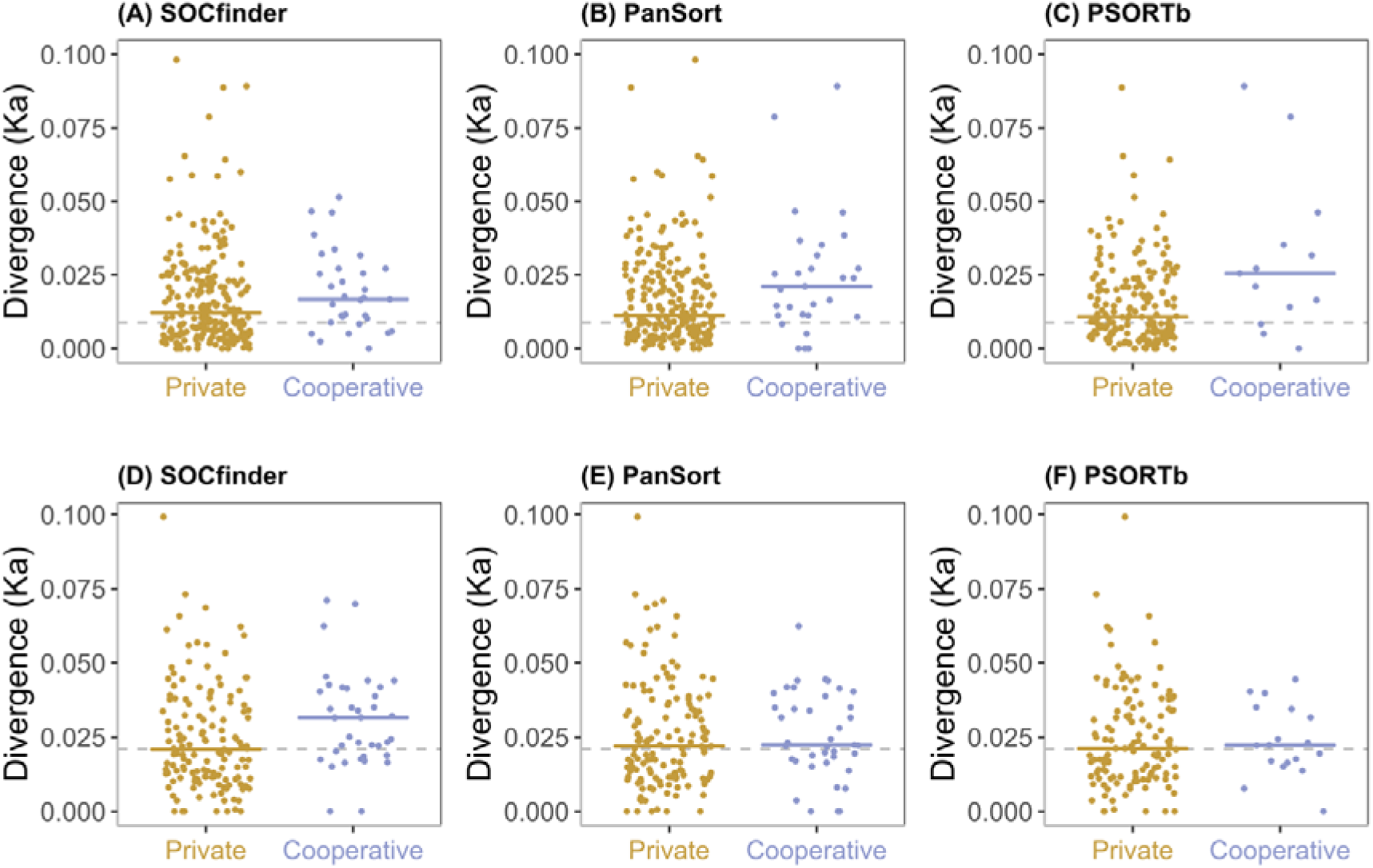
Non-synonymous divergence for private (gold) and cooperative (blue) quorum-sensing controlled genes. The top three graphs (A-C) show *P. aeruginosa*, and the bottom three graphs (D-F) show *B. subtilis*. The left graphs (A&D) show cooperative genes identified by SOCfinder. The middle graphs (B&E) show cooperative genes identified by PanSort. The right graphs (C&F) show cooperative genes identified by PSORTb. For each graph, the dotted line shows the background level of non-synonymous divergence for a set of private genes.

**Supplementary Figure 2:**
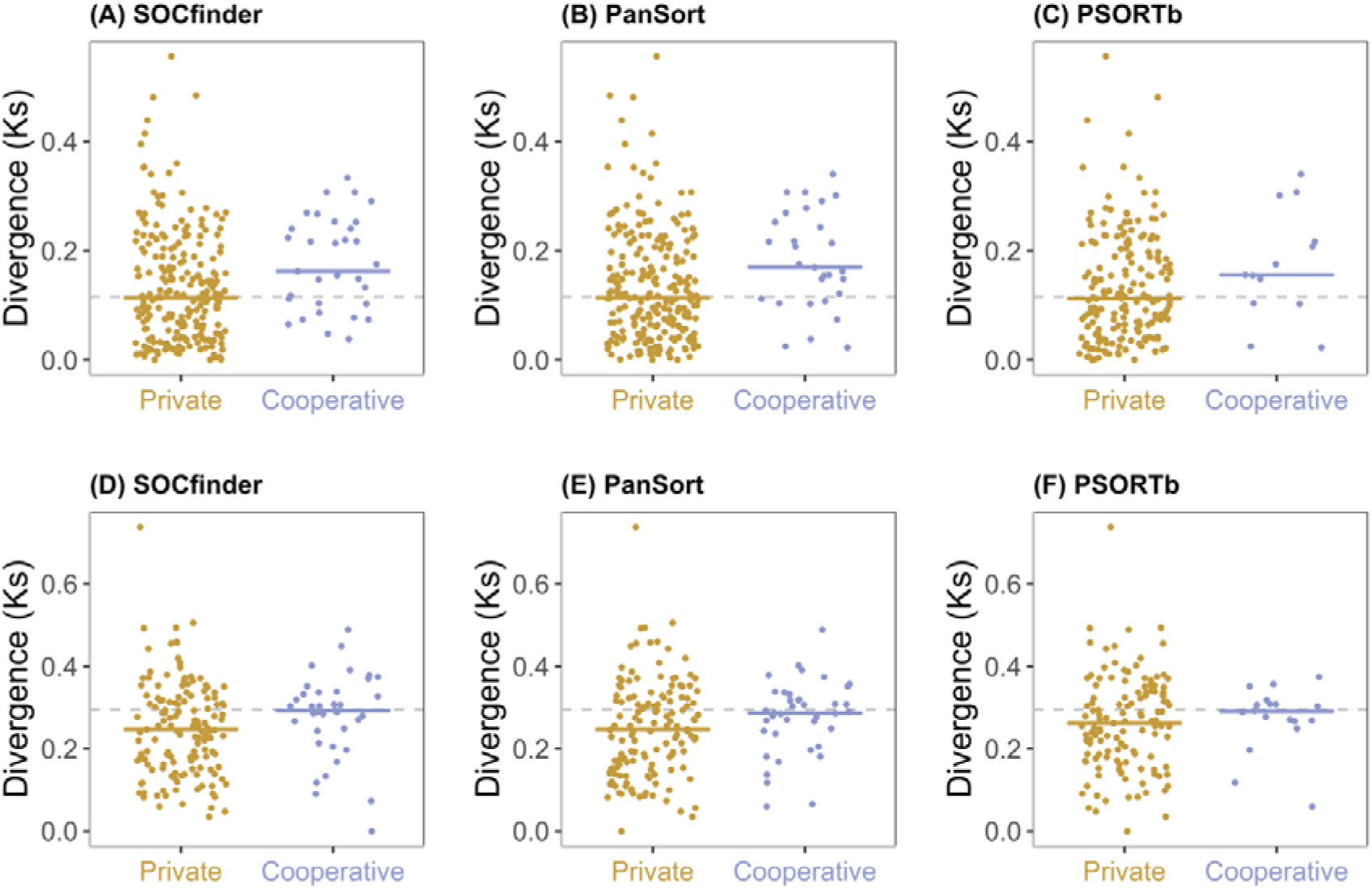
Synonymous divergence for private (gold) and cooperative (blue) quorum-sensing controlled genes. The top three graphs (A-C) show *P. aeruginosa*, and the bottom three graphs (D-F) show *B. subtilis*. The left graphs (A&D) show cooperative genes identified by SOCfinder. The middle graphs (B&E) show cooperative genes identified by PanSort. The right graphs (C&F) show cooperative genes identified by PSORTb. For each graph, the dotted line shows the background level of synonymous divergence for a set of private genes.

**Supplementary Figure 3:**
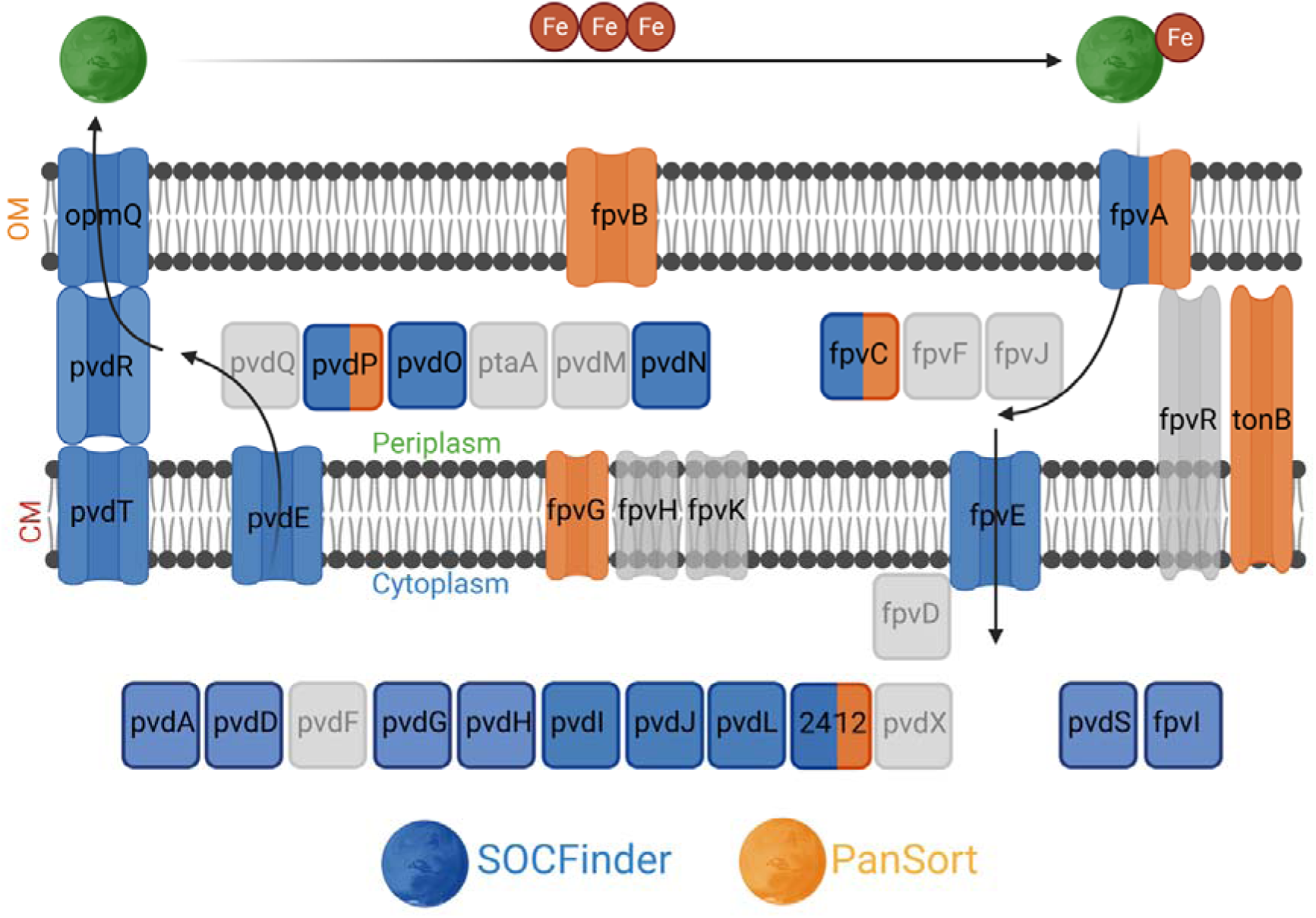
Schematic of which genes involved in the biosynthesis, export, intake, and use of pyoverdine are captured by SOCfinder and PanSort. Genes in blue are captured by SOCfinder. Genes in orange are captured by PanSort. Genes which are half blue and half orange are captured by both tools. Genes in grey are captured by no tool. Layout of genes is adapted from Ringel & Bruser (2018).

**Supplementary Figure 4:**
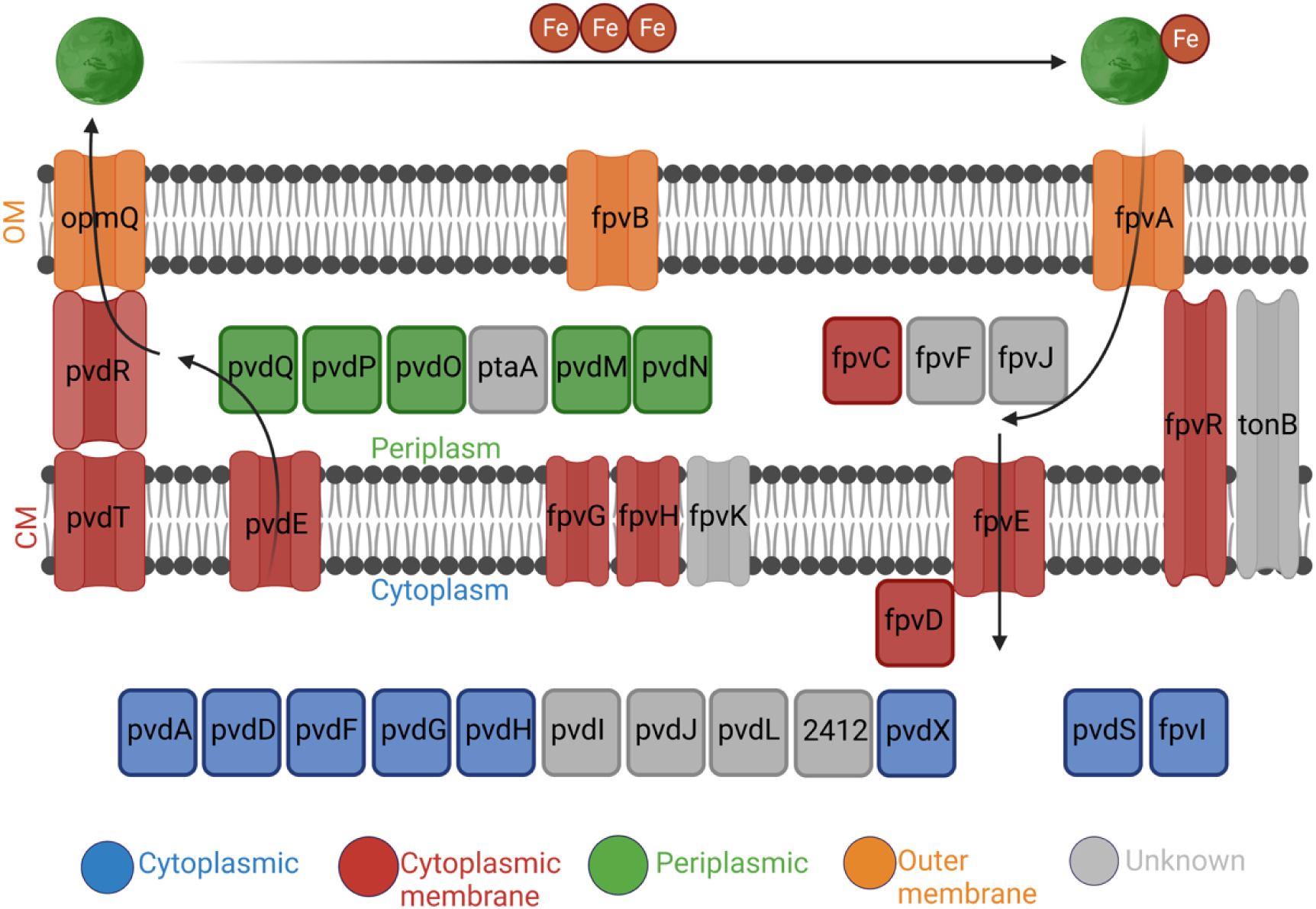
Schematic of the PSORTb subcellular localisation of proteins produced by genes involved in the biosynthesis, export, intake, and use of pyoverdine. Genes in blue code for proteins that are predicted to be cytoplasmic. Genes in red code for cytoplasmic membrane proteins. Genes in green code for periplasmic proteins. Genes in orange code for outer membrane proteins. Genes in grey code for proteins of unknown localisation. No genes code for extracellular proteins. Layout of genes is adapted from Ringel & Bruser (2018).

